# Nuclear translocation of phosphorylated YB-1 via small extracellular vesicles contributes to the malignant phenotype of triple negative breast cancer

**DOI:** 10.64898/2026.07.14.738446

**Authors:** Mark F. Santos, Yeaji Kim, Zhimin Feng, Tyler Biebighauser, Aurelio Lorico, Khalid Sossey-Alaoui

**Author notes:** Correspondence to be addressed to: Aurelio Lorico and Khalid Sossey-Alaoui. co-first authors (Mark F. Santos, Yeaji Kim). co-senior authors (Aurelio Lorico, Khalid Sossey-Alaoui).

## Abstract

Despite continuous progress in diagnosis and therapy, breast carcinoma (BC) remains a major health problem. Triple-negative (Estrogen Receptor-/Progesterone Receptor-/HER2-) breast cancer (TNBC) is the most aggressive subtype due to its high metastatic potential and resistance to chemotherapy. The Y-box binding protein 1 (YB-1) transcription factor, a protein present in both cytoplasm and nucleus, is a driver of TNBC malignancy as it stimulates its cancer stem cell phenotype and disrupts cell cycle progression. Here, we hypothesized that YB-1-containing sEVs deliver YB-1 to the nuclear compartment of recipient cancer cells and play a major role in the activation of the metastatic process. We found a selective enrichment of YB-1 in sEVs from MDA and 4T1 cells, with ∼65% and 50% of all sEVs positive for YB-1 by d-STORM. Administration of sEVs from wild-type MDA and 4T1 to their YB-1 knockout counterparts resulted in nuclear translocation of sEV-associated YB-1 and increased tumorsphere formation. Pharmacological blockade of the nuclear transport machinery based on the inhibition of the formation of the “VOR” complex (<u>V</u>AP-A-<u>O</u>RP3-<u>R</u>ab7) by PRR851 impaired both nuclear translocation and the YB-1-induced increase in tumorsphere formation. YB-1 phosphorylation at S102 was required for nuclear localization. In fact, loss of YB-1 phosphorylation inhibited tumorsphere growth and stemness of cancer cells and YB-1-positive sEVs restored the oncogenic behavior of cancer cells expressing phospho-mutant YB-1. Moreover, PRR851 inhibited the nuclear translocation of the phosphorylated form of YB-1 and the oncogenic behavior of the TNBC cells. These data support the conclusion that the nuclear translocation of sEV-associated phosphorylated YB-1 is an important factor in the malignant behavior of TNBC and a potential therapeutic target.

## Introduction

Triple-negative (Estrogen Receptor-/Progesterone Receptor-/HER2-) breast cancer (TNBC), accounting for about 15–20% of all invasive breast cancer cases in the United States, is the most aggressive subtype due to its high metastatic potential and resistance to chemotherapy [1, 2]. The molecular mechanisms that regulate its malignant progression remain mostly unknown. We recently established the Y-box binding protein 1 (YB-1) transcription factor as a novel driver of TNBC malignancy both *in vitro* and in animal cancer models [3]. This action is mediated by the stimulation of the cancer stem cell phenotype (Nanog, Oct and Sox) and disruption of cell cycle progression (Cyclin D/CDK4/6 complex and retinoblastoma protein) [4]. Accordingly, elevated levels of YB-1 are linked to poor prognosis [3].

Over the past decade, small extracellular vesicles (sEVs) have come to the limelight as pivotal vehicles of intercellular communication. They are 30-200 nanometer membrane vesicles derived by multi-vesicular bodies (exosomes) or shed from the plasma membrane (ectosomes) [5]. While present in the circulation at very high concentrations, ranging from hundreds of millions to billions per mL [5], their number increases in cancer and their cargo acquires cancer-specific, active macromolecules that facilitate metastatic dissemination. In the context of TNBC, sEVs promote invasiveness, motility, and induce surrounding stromal/immune cells in the tumor microenvironment (TME) to support TNBC growth. The same occurs at distance, in the pre-metastatic niche. Also, sEVs secreted by cancer-associated fibroblasts have been shown to drive cancer progression by transferring bioactive cargo back to TNBC cells.

As expected by its function as transcription factor, nuclear expression of YB-1 was reported in several studies [6, 7]. We previously discovered a novel intra-cellular pathway that allows sEVs to transfer their cargo into the nuclear compartment of recipient cells, where they alter gene expression [8–10]. This process is usurped by viral components of HIV-1 and cytomegalovirus [11, 12]. Upon entry into recipient target cells by endocytosis, sEVs are carried by late endosomes, defined by Rab7 GTPase, into type II nuclear envelope invaginations (NEIs), composed of both inner and outer nuclear membranes. Their entry into NEIs and docking to the outer nuclear membrane are mediated by the VOR complex, made of three proteins: ORP3 bridges VAP-A on the outer nuclear membrane with Rab7 on late endosomes [8]. The VOR complex itself is responsible for inducing the formation of new type II NEIs and for the docking of sEV-containing late endosomes near nuclear pores, by which cargoes are then transferred to the nucleoplasm [8–10].

Here, we hypothesized that YB-1-containing sEVs deliver YB-1 to the nuclear compartment of recipient cancer cells and play a major role in the activation of the metastatic process. We employed sEVs released by two well-studied models of TNBC, human MDA-MB-231 (MDA) and murine 4T1 cells, to investigate the pro-tumorigenic function of the EV-carried YB-1 protein and its relationship with the sub-cellular distribution, specifically its nuclear translocation.

We now report that YB-1 is enriched in sEVs derived from breast cancer cells, which drives their oncogenic behavior. Pharmacological blockade of the “spathasome” nuclear transport machinery[8–11, 13] inhibits the nuclear translocation of EV-associated phosphorylated YB-1 to YB-1 KO cells and the YB-1-induced malignant behavior of TNBC cells.

## Materials and Methods

### Cell Culture

The human MDA-MB-231 (MDA) and murine 4T1 triple-negative breast cancer cell lines were obtained from ATCC and cultured in RPMI-1640 medium (#10-041-CV, Corning Inc.) supplemented with 10% fetal bovine serum (FBS; Thermo Fisher Scientific), 2 mM L-glutamine, and antibiotics (100 U/mL penicillin and 100 μg/mL streptomycin). Cells were maintained at 37°C in a humidified atmosphere containing 5% CO2.

YB-1-knockout (YB-1-KO) cell lines were generated using CRISPR/Cas9-mediated gene editing as previously described [3]. Briefly, cells were transfected/electroporated with Cas9 and guide RNAs targeting the human or murine YB-1 gene. murine sgRNAs: A*C*G*GGCAGCGGCGCGGGUAG, G*G*C*GGGGACAAGAAGGUCAU and G*G*C*CCGAGCCACGGACUACG human sgRNAs: U*U*U*UCCAGCAACGAAGGUUU and U*U*C*AUCAACAGGUGAGCUGC (Synthego). Knockout efficiency was confirmed by immunoblotting for YB-1 expression.

For reconstitution studies, YB-1-KO MDA cells were stably transfected with plasmids encoding wildtype YB-1 (WT), phospho-mutant YB-1 (S102A), or phospho-mimetic YB-1 (S102D). Stable cell populations were selected using Geneticin (Thermo Fisher Scientific) and expression was verified by immunoblot analysis.

### PRR851 Treatment

PRR851 was synthesized as previously described [9, 14]. A stock solution of PRR851 was prepared in dimethyl sulfoxide (DMSO). Working solutions were prepared immediately before use by dilution into cell culture medium. Cells were treated with PRR851 at the concentrations and durations indicated in each experiment. Control cells received equivalent concentrations of DMSO.

### Isolation and Characterization of Small Extracellular Vesicles (sEVs)

Small extracellular vesicles (sEVs) were isolated from conditioned media of parental and YB-1-KO MDA and 4T1 cells. Cells were seeded at a density of 2.5 × 10^5^ cells per well in 6-well plates coated with 20 μg/mL poly(2-hydroxyethyl methacrylate) (poly-HEMA; Sigma-Aldrich) to prevent cell attachment, as previously described [15]. Cells were maintained for 72 h in serum-free RPMI-1640 medium supplemented with 2% B-27 supplement (Thermo Fisher Scientific).

At the time of collection, cell viability exceeded 90% as determined by trypan blue exclusion. Conditioned media were sequentially centrifuged at 300 × g for 10 min and 1200 × g for 20 min to remove cells and debris, followed by centrifugation at 10,000 × g for 30 min at 4 °C. The resulting supernatants were ultracentrifuged at 100,000 × g for 60 min at 4 °C using a Beckman Coulter Optima ultracentrifuge equipped with an SW41Ti rotor. Pelleted sEVs were washed once with PBS and centrifuged again at 100,000 × g for 60 min. The final pellet was passed through 0.22 μm low-protein binding filters, resuspended in sterile PBS and stored at −80 °C until use.

The size distribution and concentration of sEVs were determined by nanoparticle tracking analysis (NTA) using a ZetaView instrument (Particle Metrix GmbH). Data acquisition was performed in scatter mode using a 488 nm laser, with videos recorded at 11 positions per sample. Particle size and concentration were analyzed using the manufacturer’s software. sEV identity and purity were further evaluated by immunoblotting for the EV markers CD9 and CD81 and the negative EV marker calnexin, in accordance with MISEV 2023 guidelines [17].

### Direct Stochastic Optical Reconstruction Microscopy (d-STORM)

sEVs were immobilized on microfluidic chips supplied with the EV Profiler Kit (Oxford Nanoimaging) and processed according to the manufacturer’s instructions. Vesicles were fixed using F1 fixation solution for 10 min at room temperature. For intra-vesicular detection of YB-1, sEVs were permeabilized with 0.01% Triton X-100 in PBS for 10 min.

sEVs were immunolabeled with antibodies against YB-1 (Abcam, ab76149) together with pan-tetraspanin markers (CD9, CD63, and CD81). Antibodies were diluted in permeabilization buffer containing 1% bovine serum albumin (BSA) and 0.01% Triton X-100. Following incubation and washing, samples were imaged in freshly prepared d-STORM imaging buffer.

Single-molecule fluorescence images were acquired using a Nanoimager S Mark II microscope (Oxford Nanoimaging) equipped with a 100× oil immersion objective. Multichannel registration was calibrated using TetraSpeck fluorescent beads. Raw data were processed using NimOS software and analyzed using CODI (Oxford Nanoimaging). A minimum of 3500 individual sEVs were analyzed per condition. The percentage of YB-1-positive sEVs was calculated as the proportion of pan-tetraspanin-positive vesicles that also contained YB-1 signal.

### Western Blot

Whole-cell lysates were prepared using an ice-cold lysis buffer containing 1% Triton X-100, 150 mM NaCl, and 50 mM Tris-HCl (pH 8.0) supplemented with protease and phosphatase inhibitors. Lysates were incubated on ice for 30 min and centrifuged at 10,000 × g for 5 min at 4 °C. The resulting supernatants were collected and mixed with SDS sample buffer. Purified sEVs were lysed directly in SDS sample buffer and processed in parallel with cell lysates.

Proteins were separated by SDS-PAGE and transferred onto nitrocellulose membranes. Membranes were blocked in PBS containing 1% BSA and incubated overnight at 4 °C with primary antibodies against YB-1(Abcam, ab76149), phospho-YB-1 (S102) (Abcam, ab138654), CD9 (Santa Cruz Biotechnology, SC-20048) CD81(Invitrogen, MA5-13548), calnexin (Invitrogen, MA3-027), Lamin B1 (Cell Signaling Technology, 13435), α-tubulin (Cell Signaling Technology, 3873), β-actin (Millipore, A2228), and other indicated proteins. After washing, membranes were incubated with the appropriate secondary antibodies and visualized and analyzed using iBright FL1000 or ChemiDoc MP Imaging System (Bio-Rad).

## Immunofluorescence and Confocal Microscopy

Cells were seeded onto 8-well chamber slides (Ibidi) and allowed to adhere overnight. For sEV uptake experiments, recipient cells were incubated with purified sEVs for 5 h at 37 °C unless otherwise indicated. Cells were washed with PBS, fixed with 4% paraformaldehyde for 15 min at room temperature (RT), permeabilized using 0.2% Tween-20 in PBS for 15 min, and blocked with 1% BSA for 30 min. Samples were incubated with primary antibodies against YB-1(Abcam, ab76149), phospho-YB-1 (S102) (Abcam, ab138654) or VAP-A. Following washing, cells were incubated with species-appropriate fluorescent secondary antibodies and nuclei were counterstained with DAPI. Images were acquired using the Nanoimager or Leica DM5500 confocal microscope. Image processing and quantification were performed using Fiji/ImageJ software.

For experiments assessing nuclear localization, fluorescence intensity within DAPI-defined nuclear regions was quantified. For EV-transfer experiments, nuclear YB-1 puncta and YB-1-positive type II nuclear envelope invaginations (NEIs) were quantified as described in the figure legends.

### Nuclear and Cytoplasmic Fractionation

Nuclear and cytoplasmic fractions were prepared using NE-PER Nuclear and Cytoplasmic Extraction Reagents (Thermo Scientific, 78833). Briefly, cells were harvested and processed according to the manufacturer’s instructions/protocol to separate nuclear and cytoplasmic protein fractions. Protein concentrations were determined using DC Protein Assay Kit (Bio-Rad). Equal amounts of protein were analyzed by immunoblotting. Lamin B1 and α-tubulin were used as markers for nuclear and cytoplasmic fractions, respectively.

### Tumorsphere Formation Assay

Tumorsphere assays were performed as previously described [15, 16]. Cells were seeded into ultra-low attachment plates at a density of 4000 cells for MDA-MB-231 and 1000 cells for 4T1 per well in Spheroid media (F-12 media, B-27 supplement, EGF, bFGF and Insulin). Where indicated, purified sEVs isolated from parental or YB-1-KO cells were added at a concentration of ∼7.5 x 10^8^ particle/ml. PRR851 or vehicle control (DMSO) was added at the indicated concentrations. Tumorspheres were cultured for 12 days and imaged using Nikon Eclipse light microscope. Tumorsphere size was quantified using ImageJ software.

### Limiting Dilution Assay

Limiting dilution assays were performed to assess stem cell frequency. Cells were seeded into ultra-low attachment plates at serial dilutions ranging from 1024 to 8 cells for MDA-MB-231 and 350 to 3 cells for 4T1 cells per well in 96-well plate. Following incubation for 14 days, wells were scored for tumorsphere formation. Stemness frequency was calculated using the ELDA Analysis Software as described (http://bioinf.wehi.edu.au/software/elda/index.html).

### Colony Formation Assay

Cells were seeded into 6-well plate at a density of 3000 cells for MDA-MB-231 and 1000 cells for 4T1 cells per well and cultured in complete medium in the presence of vehicle control (DMSO) or PRR851. After culturing for 10 days, colonies were fixed using 4% paraformaldehyde (PFA) at room temperature for 20 minutes and stained with 0.25% crystal violet solution. Colonies were imaged with ChemiDoc Imaging System (Bio-Rad) and quantified using ImageJ software.

### Cell Viability Assay

Cell viability was assessed using the MTT assay. Cells were seeded into 96-well plate [Corning] and treated with vehicle control (DMSO) or PRR851 for the indicated time periods. MTT reagent was added according to the manufacturer’s instructions. Absorbance was measured using a microplate reader at 560 nm, and viability was expressed relative to vehicle-treated controls.

### Statistical Analysis

All experiments were performed using at least three independent biological replicates unless otherwise indicated. Data are presented as mean ± standard deviation (SD). Statistical analyses were performed using GraphPad Prism version 10.1.1. Comparisons between two groups were performed using an unpaired two-tailed Student’s t-test. Comparisons involving multiple groups were analyzed using one-way ANOVA. Differences were considered statistically significant at p < 0.05.

## Results

### sEVs derived from breast cancer cells are enriched in YB-1, which drives their malignant behavior

Interrogation of the Vesiclepedia database (http://microvesicles.org/) revealed the presence of YB-1 in sEVs of several BC cell lines, including MDA and 4T1 TNBC cells as well as of stromal (fibroblasts) and immune (T cells and dendritic) cells (Suppl. Table 1). To explore the potential role of TNBC EV-associated YB-1, we first isolated sEVs from MDA and 4T1 BC cells using a standard ultracentrifugation-based protocol. To ensure that sEVs were produced independently of cell attachment, the cultures were maintained in serum-free media on poly-HEMA-coated plates. Nanoparticle tracking analysis (NTA), direct stochastic optical reconstruction microscopy (d-STORM) and immunoblotting for established EV markers were used to confirm particle identity and purity, according to MISEV guidelines [17] (Fig. 1A-D). We then assessed by NTA [15, 18] whether the genetic deletion of YB-1 affects sEV secretion. The size distribution and concentration of sEVs from parental MDA and 4T1 cells and their YB-1-knockout (YB-1-KO) counterparts were comparable across both cell lines (Figure 1A). For MDA cells, the average EV concentration was 1.22 ± 0.17 × 10^10^ particles/mL for parental cells and 1.08 ± 0.15 × 10^10^ particles/mL for YB-1-KO cells (p = 0.17), with corresponding sizes of 153.8 ± 53.5 nm and 158.0 ± 62.6 nm, respectively (p = 0.90). For 4T1 cells, the average EV concentration was 1.75 ± 0.37 × 10^10^ particles/mL for parental cells and 2.33 ± 0.66 × 10^10^ particles/mL for YB-1-KO cells (p = 0.10), with corresponding sizes of 153.1 ± 60.4 nm and 165.2 ± 69.8 nm, respectively (p = 0.75). These results indicate no statistically significant difference in overall EV yield or size distribution. The lack of changes suggests that YB-1 is not required for bulk EV production or release, and that YB-1-KO does not substantially impact EV biogenesis under the conditions tested.

**Fig. 1.**
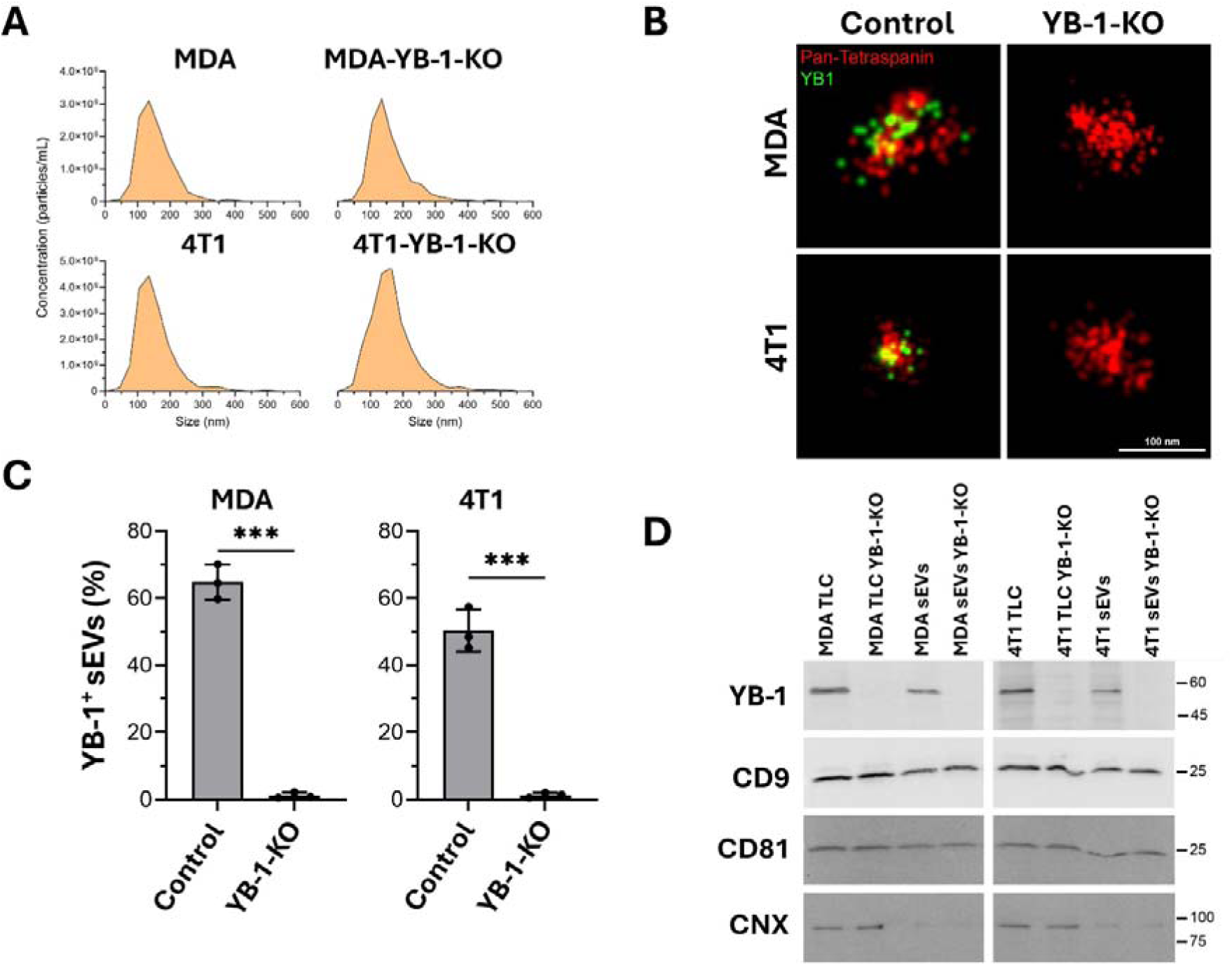
Presence of YB-1 in sEVs from TNBC cells. **(A)** Nanoparticle tracking analysis and **(B)** representative d-STORM images of sEVs from parental and YB-1-KO TNBC cells. **(C)** Percentage of YB-1-positive sEVs from parental and YB-1-KO TNBC cells as imaged by d-STORM. Data represent mean ± SD from at least three independent experiments (***, p < 0.001). **(D)** Parental MDA and 4T1 cells, their YB-1-KO derivatives and their respective sEVs were analyzed by immunoblotting for YB-1, CD9, CD81 and calnexin (CNX; negative EV marker).

We next asked whether YB-1 is physically incorporated into sEVs. Since YB-1 is not a membrane protein, we permeabilized the EVs employing a method developed by our team for intra-vesicular proteins [19]. Using d-STORM, we observed a strong co-localization of YB-1 with pan-tetraspanin markers in sEVs derived from parental MDA and 4T1 cells (Figure 1B). In contrast, sEVs from YB-1-KO cells showed virtually no YB-1 signal. Quantification revealed that approximately 65% and 50% of sEVs from MDA and 4T1 parental cells, respectively, were YB-1-positive (Figure 1C). This indicates that YB-1 is selectively enriched in sEVs.

We confirmed these findings by immunoblot analysis of both whole-cell lysates and purified EV fractions (Figure 1D). YB-1 was readily detectable in sEVs from both MDA and 4T1 control cells but absent in sEVs from YB-1-KO cells. The canonical EV markers CD9 and CD81 were present in all the samples, while calnexin, a marker typically excluded from sEVs, was undetectable, confirming the purity of sEV preparations.

To assess for the biological effects of TNBC cell-derived YB-1^+^ sEVs, we employed a 3D-tumorsphere growth assay as a readout of oncogenic behavior, and showed that the growth potential of the tumorspheres derived from MDA (Figure 2A) or 4T1 cells (Figure 2B) in serum-free medium was significantly enhanced upon cell exposure to sEVs derived from parental cells (YB-1^+^ sEVs), compared to cells exposed to sEVs derived from the YB-1-KO cells (YB-1^-^sEVs). No significant difference in tumorsphere growth was observed between cells treated with YB-1^+^ sEVs and those treated with complete growth media (Complete Media), that contains sEVs. Importantly, we observed a significant increase in growth of tumorspheres of YB-1-KO-MDA cells (Figure 2C) or YB-1-KO-4T1 cells (Figure 2D) treated with YB-1^+^ sEVs, compared to those treated with YB-1^-^sEVs. Thus, we confirmed the effect of YB-1-enriched sEVs in the activation of the oncogenic behavior of BC cells.

**Fig. 2.**
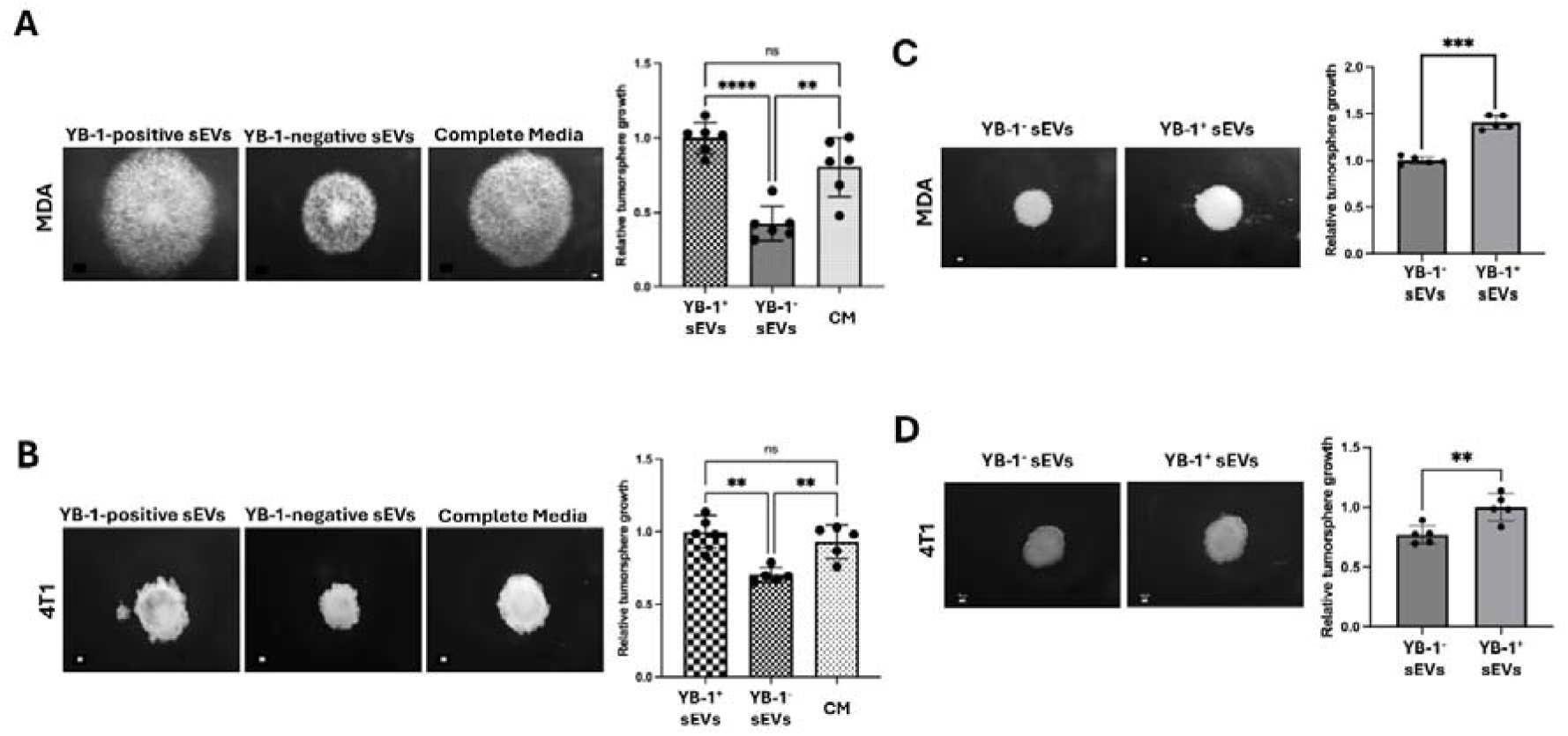
Effect of sEV-associated YB-1 on tumorsphere growth. 3D tumor spheroid growth assays using MDA **(A)**, 4T1 **(B)** and their respective YB-1-knockout cells **(C-D)**. Cells in serum-free medium were supplemented with either purified sEVs from parental (YB-1-positive sEVs) or YB-1 knockout cells (YB-1-negative sEVs). Cells grown in serum-supplemented medium (complete) without addition of sEVs were used as control. In the right panels the relative tumorsphere growth is presented (means ± S.D. with the individual value of each experiment (*n* = 5-6). Scale bars, 100 µm. ns, not significant; *, p < 0.05; **, p < 0.01; ***, p < 0.001; ANOVA.

### YB-1 phosphorylation at S102 is required for its nuclear localization

YB-1 phosphorylation at serine residue 102 (S102, Figure 3A) has been associated with the enhanced oncogenic activity of YB-1 [20–23]. By immunoblotting of whole-cell lysates of cells serum-starved (Starved) or grown in complete growth media (CM) using an anti-YB-1 Ab specific for the S102 phosphorylated form (pYB-1(S102)), we showed that levels of pYB-1(S102) were 4-(4T1) and 10-fold (MDA) increased in cells grown in CM compared to starved cells (Figure 3B). Next, we stably expressed either wildtype YB-1 (WT), phospho-mimetic YB-1 (S102D) or phosphor-mutant YB-1 (S102A) in YB-1-KO-MDA cells. pYB-1 levels were significantly diminished in cells expressing phospho-mutant S102A YB-1, compared to cells expressing either WT or phospho-mimetic S102D YB-1 (Figure 3C). Since phosphorylation of YB-1 is required for its translocation to the nucleus, where it acts as oncogenic transcription factor [23], immunofluorescence revealed enrichment of pYB-1-S102 in the nuclei of MDA cells (Figure 3D). We validated these findings by performing an immunoblot analysis on cytoplasmic and nuclear cell lysate fractions and found a 10-fold increase of YB-1 in the nuclear fraction of MDA cells grown in the presence of complete media vs. cells serum-starved (Figure 3E). Immunofluorescence using YB-1-specific antibody (Figure 3F) also confirmed the nuclear enrichment of the phospho-mimetic S102D form of YB-1 in the nuclei of MDA cells compared to the phosphor-mutant S102A form of YB-1 as quantitated in Figure 3G.

**Fig. 3.**
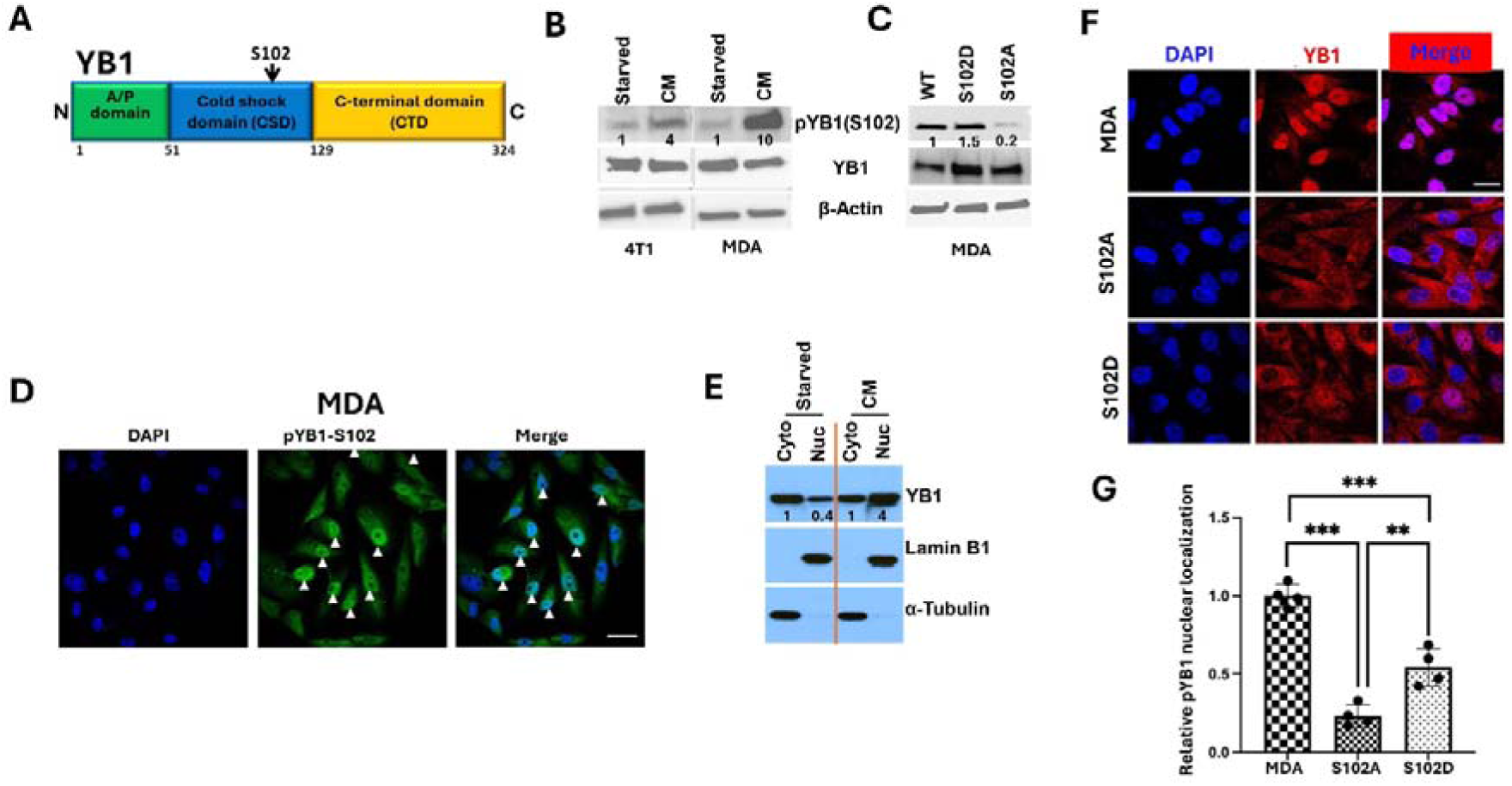
YB-1 phosphorylation at S102 is required for its nuclear localization. **(A)** Domain structure of YB-1 protein. The location of Serine 102 (S102) is indicated by the black arrow. **(B)** Representative immunoblot of total cell lysates from 4T1 and MDA cells (serum-starved or in complete medium (CM)) probed with anti-YB-1 or anti-pYB-1(S102) antibodies. β-actin was used as loading control. **(C)** Representative immunoblot of total cell lysates from YB-1-KO-MDA cells stably transfected with wildtype YB-1 (WT), phospho-mimetic YB-1 (S102D) or phospho-mutant YB-1 (S102A) and probed with either anti-YB-1 or anti-pYB-1(S102) antibodies. β-Actin was used as loading control. **(D)** Representative confocal microscopy images of MDA cells stained with anti-pYB-1 antibodies (red). Cell nuclei were counterstained with DAPI (blue). The white arrows point to the intense staining of YB-1 in the nucleus. **(E)** Representative immunoblot of cytoplasmic and nuclear fractions of MDA cells (serum-starved or in complete medium (CM)) probed with antibodies against YB-1, Lamin B1 (nuclear fraction marker) or α-Tubulin (cytoplasmic fraction marker). The numbers under each band represent the fold change in signal intensity with respect to its respective control after normalization for the loading control signal. Data shown are representative of 3 independent experiments. **(F)** Representative confocal microscopy images of YB-1-KO MDA with or without stable expression of phospho-mutant (S10A) YB-1 or phospho-mimetic (S102D) YB-1 after staining with anti-pYB-1 antibodies (red). Cell nuclei were counterstained with DAPI (blue). **(G)** Relative pYB-1 nuclear localization for phospho-mimetic YB-1 (S102D) or phospho-mutant YB-1 (S102A) (mean ± S.D. with the individual value of each experiment (*n* = 4). Data are the mean ± SD (**p < 0.01, ***p < 0.001, ANOVA).

### Loss of YB-1 phosphorylation inhibits 3D tumorsphere growth and stemness of cancer cells

To assess the effect of loss of phosphorylation of YB-1 on its oncogenic activity, we performed 3D-tumorsphere growth assays (Figure 4A) and showed that expression of the phospho-mutant S102A YB-1 in YB-1-KO cells resulted in inhibition of MDA tumorsphere growth to levels similar to those exhibited by YB-1-KO cells. Tumorsphere growth was partially restored upon expression of phospho-mimetic S102D YB-1(Figure 4B). The reason for the only partial restoration of tumorsphere growth in the S102D-expressing cells could be attributed to the lower levels of expression of S102D in MDA cells compared to endogenous YB-1. Nonetheless, the data show that phosphorylated YB-1 and its nuclear localization are required for the activation of the oncogenic behavior of BC cells. These findings were further confirmed using the limiting dilution assay (Figure 4C) and showed that YB-1-KO and expression of S102A decreased MDA stemness (Figure 4D), and stemness was partially restored in YB-1-KO MDA cells expressing S102D YB-1.

**Fig. 4.**
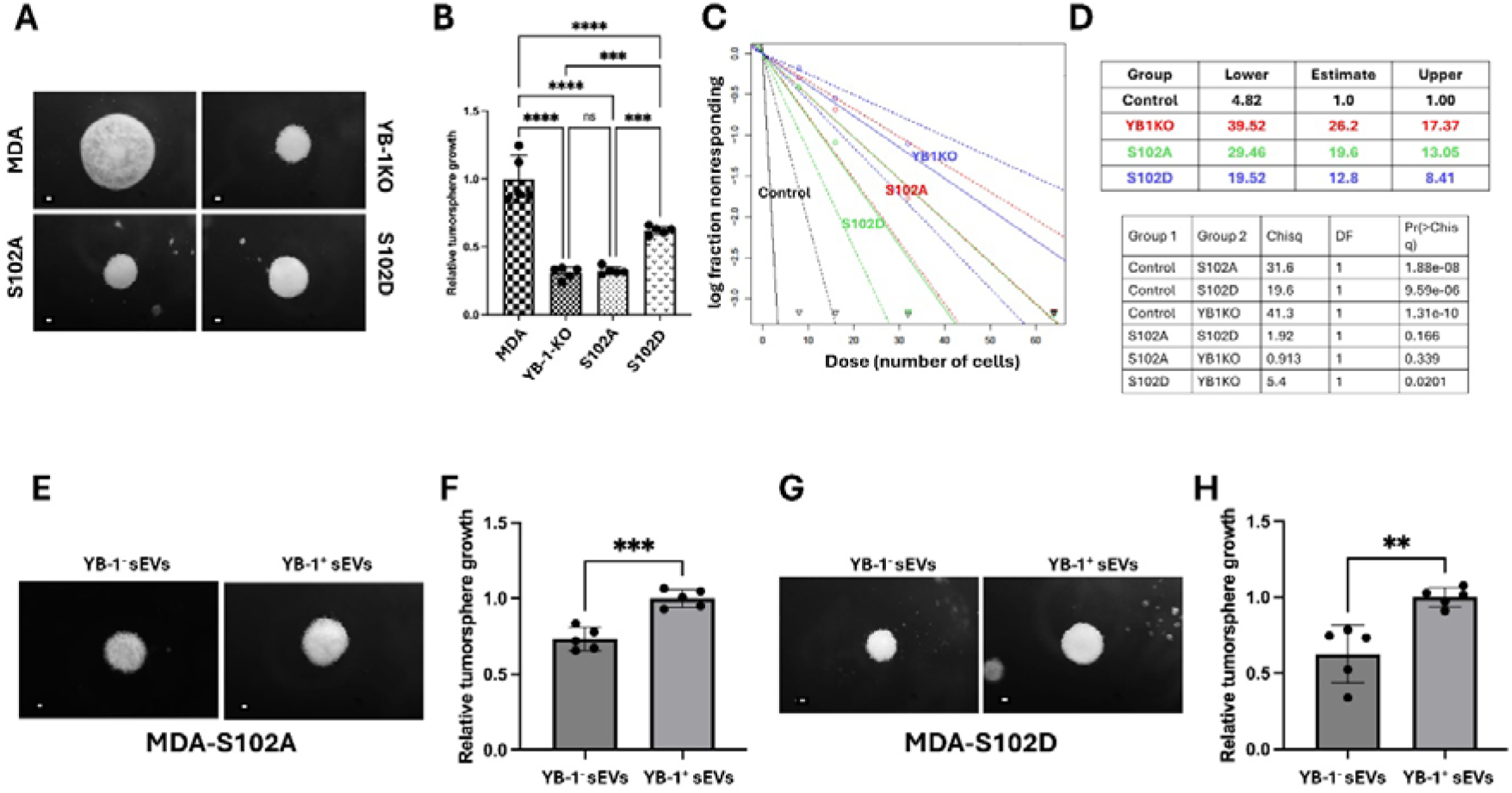
Loss of YB-1 phosphorylation inhibits 3D tumorsphere growth and stemness of cancer cells and YB-1-enriched sEVs restore the oncogenic behavior of cancer cells expressing phospho-mutant YB-1. **(A)** Representative micrographs of 3D-tumorspheres of parental MDA cells, their YB-1-knockout derivatives (YB-1KO), YB-1-KO cells with stable expression of phospho-mutant (S10A) YB-1 or phospho-mimetic (S102D) YB-1. Quantification is shown in **(B)** (mean ± S.D. with the individual value of each experiment (*n* = 5). Scale bar: 100 µm. Data are the mean ± SD (***p < 0.01; ****p < 0.001; ns, not significant, ANOVA). **(C)** Limiting dilution assay of parental MDA cells, their YB-1-knockout derivatives (YB-1KO), YB-1-KO cells with stable expression of phospho-mutant (S102A) YB-1 or phospho-mimetic (S102D) YB-1. Quantification is shown in **(D)**. Data shown are representative of 3 independent experiments. **(E-H)** 3D tumor spheroid growth assays using YB-1-KO MDA cells expressing phospho-mutant (S102A) YB-1 **(E)** or phospho-mimetic (S102D) YB-1 **(G)**. Cells in serum-free medium were supplemented with purified sEVs from parental (YB-1^+^ sEVs) or YB-1 knockout cells (YB-1^-^sEVs). Quantification is shown (**F** and **H**) (means ± S.D. with the individual value of each experiment (n = 5-6). Scale bars, 100 µm. **, p < 0.05; ***, p < 0.001; ANOVA.

### YB-1-enriched sEVs restore the oncogenic behavior of cancer cells expressing phospho-mutant YB-1

To test the effect of exogenous YB-1^+^ sEVs on tumorsphere growth of cells lacking pYB-1, MDA cells expressing phospho-mutant (S10A) YB-1, that lacks nuclear translocation ability, were supplemented with either YB-1^+^ or YB-1^-^sEVs. Cell supplemented with YB-1^+^ sEVs showed significant tumorsphere growth compared to those supplemented with YB-1^-^sEVs (Figure 4E-F). These findings were further confirmed with MDA cells expressing phospho-mimetic (S102D) YB-1, which showed a modest increase in tumorsphere growth after treatment with YB-1^+^ sEVs (Figure 4G-H). Thus, we confirmed the ability of YB-1^+^ sEVs to restore the oncogenic behavior of cells lacking pYB-1.

### Pharmacological blockade of the “spathasome” nuclear transport machinery inhibits the oncogenic behavior of cancer cells

We (the Lorico lab) recently developed a small molecule inhibitor (PRR851) that specifically targets the nuclear transport machinery named “spathasome”, consisting of elongated EV-containing late endosomes in type II nuclear envelope invaginations, part of the nucleoplasmic reticulum and, therefore, inhibits the nuclear translocation of sEV cargo [9]. Using the MTT assay, we found that treatment of MDA (Figure 5A) or 4T1 cells (Figure 5B) with 10 μM PRR851 for up to 4 days has no significant effect on cell viability, which confirms previous finding establishing PRR851 as a very safe compound [9, 14]. The effect of PRR851 on the oncogenic behavior of MDA and 4T1 cells was, however, very significant. In fact, PRR851 inhibited the colony formation potential of MDA (Figure 5C) and 4T1 cells (Figure 5D). PRR851 also inhibited 3D tumorsphere growth of MDA cells (Figure 5E) and 4T1 cells (Figure 5F). Finally, we found PRR851 to significantly inhibit the stemness of both MDA (Figure 5G) and 4T1 cells (Figure 5H). Together, these data confirm that PRR851 is a potent inhibitor of the oncogenic behavior of BC cells through its blockade of the nuclear translocation of YB-1^+^ sEVs.

**Fig. 5.**
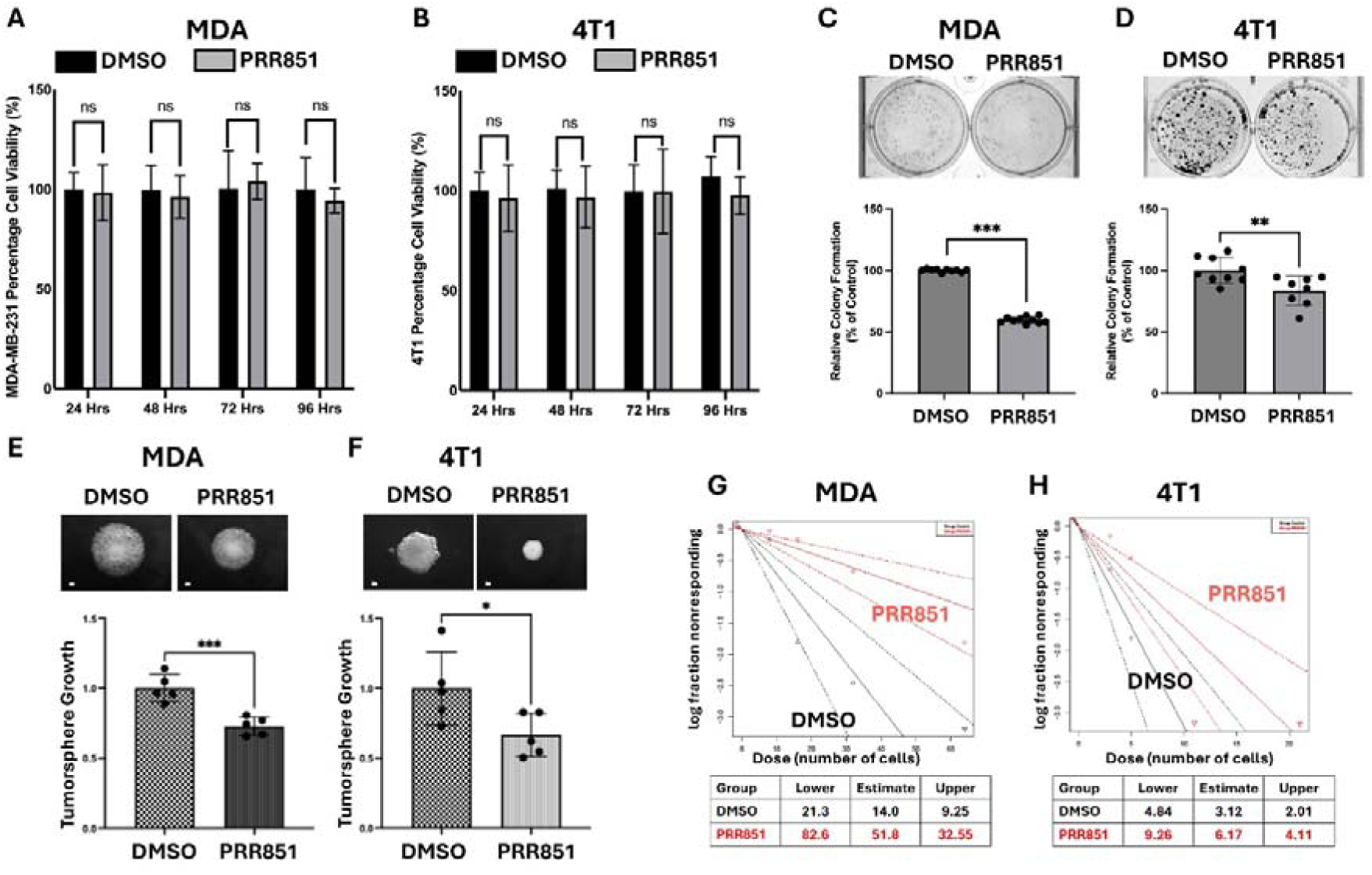
The pharmacological blockade of the “spathasome” nuclear transport machinery inhibits the oncogenic behavior of cancer cells. **(A-B)** Cell proliferation of MDA (A) and 4T1 cells (B) was measured by the MTT assay in the presence of PRR851 (10 μM) or control DMSO at the indicated time points. Data represent mean ± SD of at least 6 replicates (ns, not significant, Student t-test). **(C-D)** Colony formation assay of MDA **(C)** and 4T1 cells **(D)** in the presence of PRR851 (10 µM) or control DMSO. Data are the mean ± SD of at least 3 replicates (*p < 0.05, **p < 0.01, Student t-test). **(E-F)** Representative images of tumorspheres of MDA **(E)** or 4T1 cells **(F)** in the presence of PRR851 (10 μM) or control DMSO. Scale bar, 100 µm. **(G-H)** Limiting dilution assay of MDA **(G)** or 4T1 cells **(H)** in the presence of PRR851 (10 μM) or DMSO control. Data shown are representative of 3 independent experiments.

### PRR851 blocks the nuclear translocation of pYB-1

To assess the relevance of YB-1 phosphorylation status on the effect of PRR851-mediated effect on the oncogenic behavior of TNBC cells, we performed immunoblot analyses with anti-pYB-1 antibody on nuclear and cytosolic fractions of MDA and 4T1 cells (Figure 7A). Treatment with PRR851 significantly decreased the amount of pYB-1 in the nuclear fraction of PRR851-treated cells by ∼5 fold in MDA cells and ∼2.5 fold in 4T1 cells (Figure 6A). These findings were confirmed using immunofluorescence of PRR851-treated MDA cells that were stained with anti-pYB-1 antibody (Figure 6B). Nuclear pYB-1 was significantly inhibited in PRR851-treated cells both at 10 and 30 µM (Figure 6C). These findings clearly demonstrate that PRR851 not only blocks the nuclear transport machinery for exogenous sEVs, but also inhibits the nuclear translocation of the YB-1 protein.

**Fig. 6.**
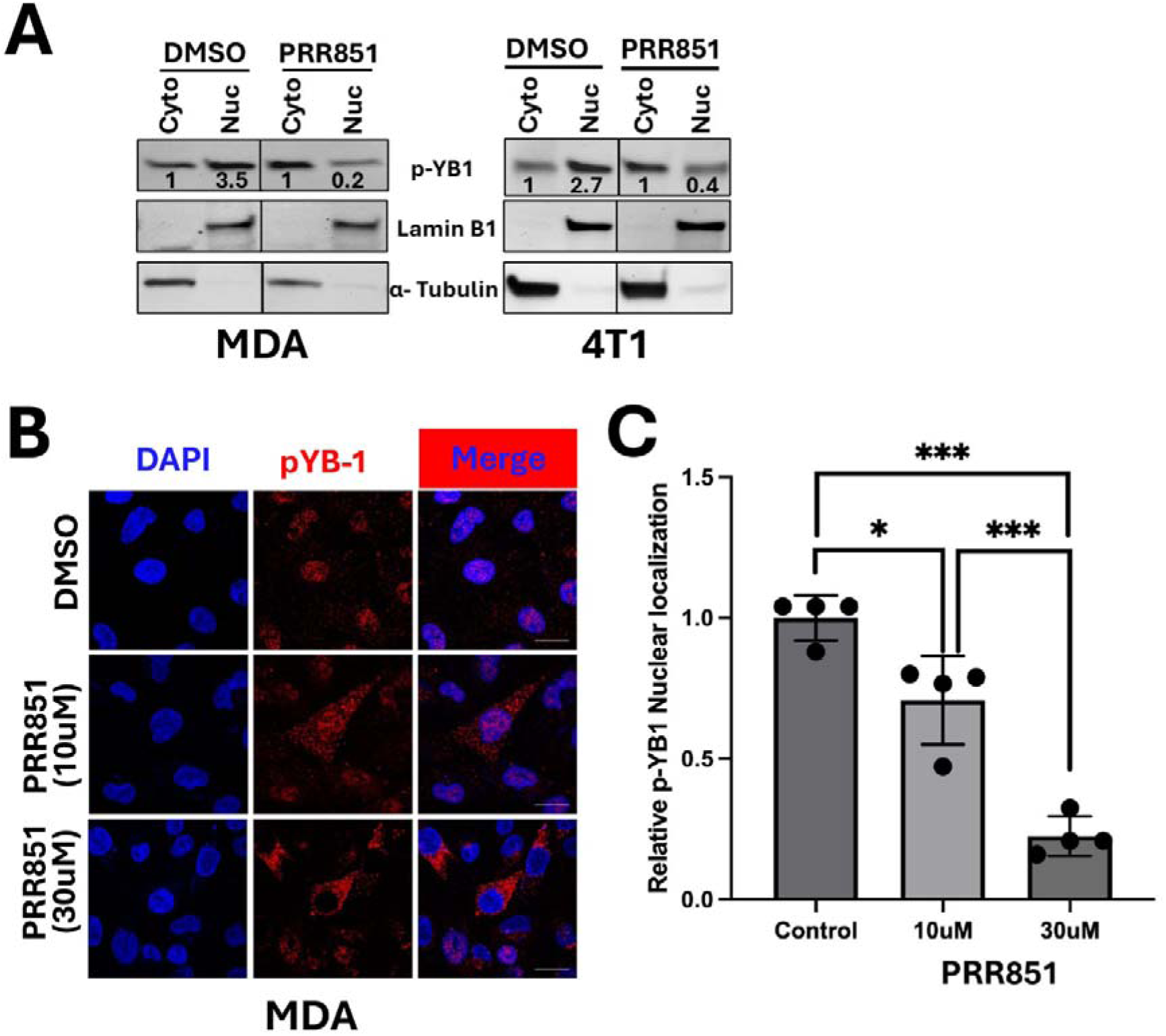
PRR851 blocks the nuclear translocation of pYB-1. **(A)** Representative immunoblots of cytoplasmic and nuclear fractions of MDA and 4T1 cells treated with either DMSO or PRR851 probed with antibodies against pYB-1, lamin B1 (nuclear fraction marker) or α-tubulin (cytoplasmic fraction marker). The numbers under each WB band represent the fold change in signal intensity with respect to control after normalization to the loading control signal. **(B)** Representative confocal microscopy images of MDA cells treated with either DMSO or PRR851 stained with anti-pYB-1 antibodies (red). Cell nuclei were counterstained with DAPI (blue). Scale bar: 20 µm. Quantification is shown in **(C)** (mean ± S.D. with the individual value of each experiment (*n* = 4) (*p < 0.05, ***p < 0.001, ANOVA).

**Fig. 7.**
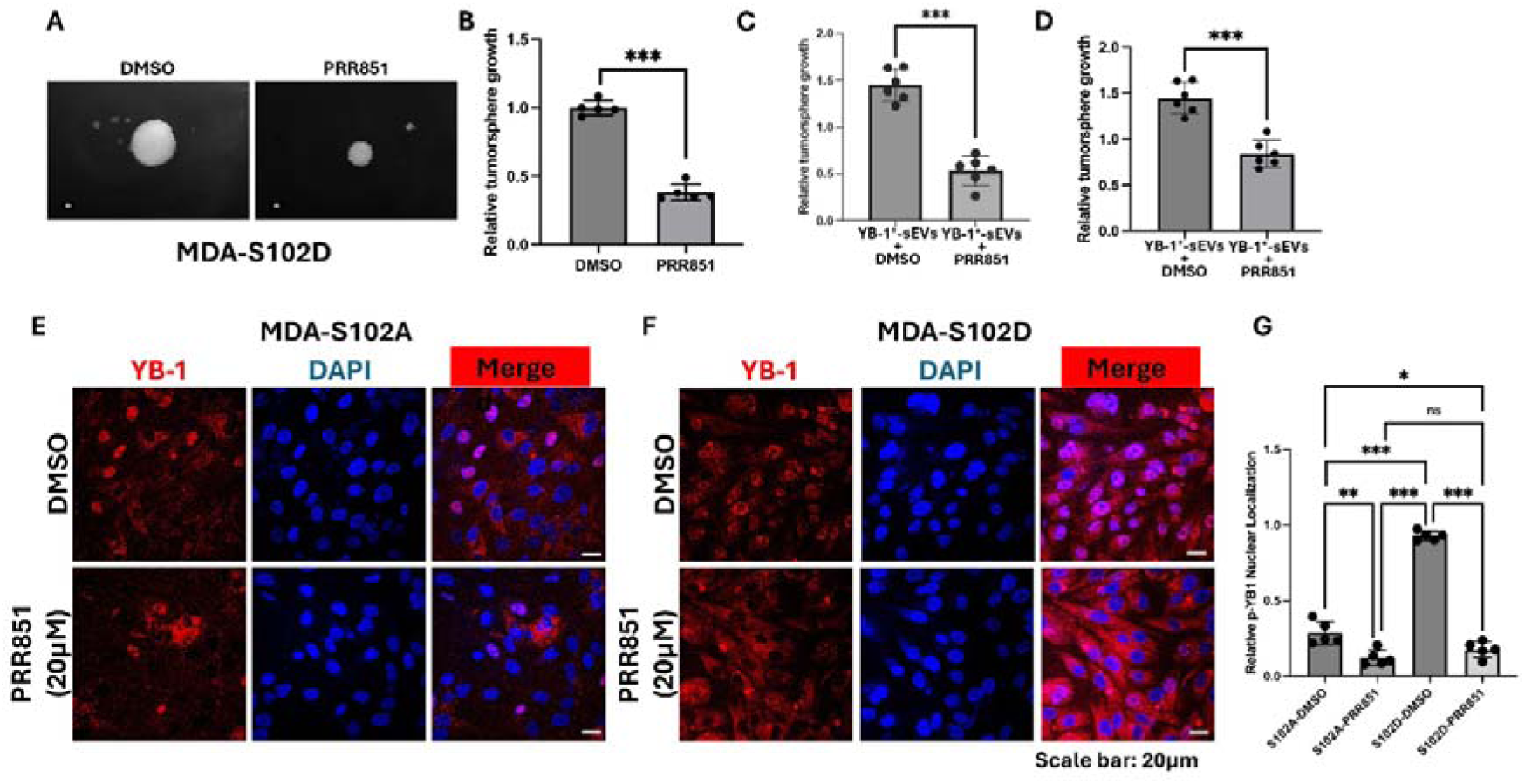
The PRR851-mediated blockade of nuclear translocation of YB-1-enriched sEVs inhibits oncogenic behavior of cancer cells. **(A)** Tumorsphere growth of S102D-expressing MDA cells in the presence of DMSO or PRR851. Scale bar: 100µm. Quantification is shown in **(B). (C-D)** Quantification of tumorsphere growth of YB-1-KO MDA cells supplemented with YB-1^+^ sEVs or YB2-sEVs and treated with PRR851 or control DMSO. **(E-F)** Confocal microscopy of YB-1-KO MDA cells expressing either phospho-mutant S102A (E) or phospho-mimetic S102D YB-1 in the presence of DMSO or PRR851, that were stained for YB-1 (Red). Nuclei were counterstained with DAPI (blue). Scale pbar: 20µm. Quantification is shown in **(G)**.

### The PRR851-mediated blockade of nuclear translocation of YB-1-enriched sEVs inhibits oncogenic behavior of cancer cells

To assess the effect of PRR851-mediated blockade on YB-1 nuclear translocation and tumorsphere growth, YB-1-KO MDA cells expressing YB-1 phospho-mimetic S102D were treated with 10 µM PRR851. Treatment with PRR851 significantly inhibited tumorsphere growth (Figure 7A-B). Next, YB-1-KO MDA (Figure 7C) or 4T1 cells (Figure 7D) grown in serum-free media were supplemented with YB-1^+^ sEVs and treated with either PRR851 or DMSO, and tumorsphere growth was assessed. Treatment with PRR851 significantly inhibited tumorsphere growth of cells supplemented with YB-1^+^ sEVs, while DMSO had no effect, suggesting that the nuclear translocation of sEVs that are enriched in YB-1 is required for tumorsphere growth. The PRR851-mediated-inhibition of YB-1 nuclear translocation was further confirmed using immunofluorescence of PRR851-treated S102A- and S102D-expressing YB-1-KO MDA cells (Figure 7E-G). Thus, we demonstrate that phosphorylation and nuclear translocation of YB-1 are required for its oncogenic activity, and that this phenotype can be inhibited by PRR851.

## Discussion

The relevance of YB-1 for the malignant phenotype of TNBC has been substantiated by experimental findings of several groups, including ours [3, 7, 24–29]. Among other pro-malignant functions, YB-1 is a crucial RNA-binding protein responsible for selectively sorting specific microRNAs (like miR-223) and other non-coding RNAs into exosomes that regulate intercellular communication in cancer and inflammation (e.g., promoting cell growth, angiogenesis or dampening inflammation) [30–32].

Here, to extend these observations to TNBC-derived sEVs carrying phosphorylated YB-1, we used a combination of state-of-the-art EV techniques (d-STORM, NTA and confocal microscopy), 3D tumorsphere growth assays, and genetic and pharmacologic targeting assays. Our observation that over 50% of sEVs released in the extracellular milieu by MDA and 4T1 cells contained YB-1, supports our hypothesis that the intercellular communication mediated by sEV-associated YB-1 is critical for TNBC malignancy. Our findings are in line with the numerous reports of pro-malignant function of many sEV-carried proteins, driving disease progression for most cancer types, including TNBC [15, 33–43].

Our findings identify YB-1-enriched sEVs as critical mediators of intercellular communication in TNBC and establish YB-1 S102 phosphorylation as a central mechanism regulating its oncogenic activity and nuclear trafficking. Together, these data support a model in which TNBC cells exploit YB-1-positive sEVs to propagate malignant signaling within the tumor microenvironment and potentially to distant organs, thereby enhancing tumor aggressiveness, stemness, and metastatic potential.

We first demonstrated that YB-1 is selectively enriched within sEVs derived from TNBC cells, while deletion of YB-1 does not significantly alter overall sEV production, size distribution, or release. These observations suggest that YB-1 is not required for bulk sEV biogenesis but instead regulates a functionally selected cargo incorporated into a subset of sEVs with specialized oncogenic functions. The enrichment of YB-1 in approximately 50– 65% of TNBC-derived sEVs further highlights the heterogeneity of the sEV compartment and raises the possibility that YB-1-positive sEVs represent a biologically distinct subpopulation dedicated to tumor-promoting functions.

Functionally, YB-1-positive sEVs strongly enhanced tumorsphere formation and restored oncogenic behavior in YB-1-deficient cells, demonstrating that extracellular YB-1 can compensate for the loss of endogenous YB-1 signaling. These findings extend previous reports implicating YB-1 in proliferation, stemness, epithelial-to-mesenchymal transition, and therapy resistance [3, 20, 23–25, 27, 30, 44–49], and suggest that TNBC cells can horizontally transfer malignant phenotypes through EV-mediated delivery of YB-1. Because tumorsphere growth and limiting dilution assays are associated with stem cell-like and tumor-initiating properties, our data further supports a role for YB-1-positive sEVs in maintaining cancer stem cell phenotypes that contribute to metastasis and recurrence.

A major mechanistic finding of this study is the requirement of YB-1 phosphorylation at S102 for its nuclear localization and oncogenic activity. Previous studies have shown that S102 phosphorylation enhances YB-1 transcriptional activity downstream of AKT/RSK signaling pathways, to enhance the oncogenic activity of YB-1 (Reviewed in [23]). Our results expand these observations by demonstrating that phospho-deficient S102A YB-1 markedly suppresses tumorsphere growth and stemness, whereas phospho-mimetic S102D YB-1 partially restores these phenotypes. Importantly, YB-1-positive sEVs were capable of rescuing oncogenic behavior even in cells expressing the S102A mutant, indicating that extracellular delivery of YB-1 can bypass deficiencies in endogenous YB-1 signaling. These findings suggest that EV-mediated YB-1 transfer may amplify oncogenic signaling networks within the tumor microenvironment and contribute to functional plasticity among TNBC cells.

Our data also reveal that YB-1-positive sEVs undergo active nuclear trafficking through the spathasome-dependent transport machinery associated with NEIs. Confocal imaging demonstrated Pharmacological inhibition of the spathasome pathway using PRR851 significantly inhibited YB-1 nuclear translocation, providing additional evidence that YB-1 cargo delivered by sEVs can access the nuclear compartment. This observation is particularly significant because it identifies an additional mechanism by which sEV cargo can directly influence nuclear signaling and transcriptional regulation in recipient cancer cells. Nuclear delivery of sEV-associated YB-1 may therefore represent a previously underappreciated mechanism of oncogenic signal propagation in TNBC.

The PRR851-mediated inhibition of the spathasome pathway using strongly suppressed tumorsphere growth, colony formation, stemness, and nuclear accumulation of phosphorylated YB-1 without significantly affecting cell viability. These findings indicate that TNBC cells are highly dependent on YB-1 nuclear trafficking for the maintenance of malignant phenotypes. Importantly, PRR851 blocked both endogenous YB-1 nuclear translocation and the nuclear delivery of YB-1-positive sEV cargo, highlighting the central role of nuclear EV trafficking in TNBC progression. Because PRR851 selectively impairs oncogenic behavior without overt cytotoxicity, targeting nuclear EV transport pathways may represent a promising therapeutic strategy with potentially reduced systemic toxicity compared to conventional chemotherapy.

Collectively, these findings support a model in which TNBC cells establish a feed-forward oncogenic communication network through secretion of YB-1-enriched sEVs. In this model, extracellular YB-1 is transferred to recipient cells, trafficked through the spathasome-associated nuclear transport pathway, and accumulates in the nucleus in a phosphorylation-dependent manner to promote transcriptional programs linked to stemness, survival, and tumor progression. Such mechanisms may contribute not only to intra-tumoral heterogeneity but also to metastatic niche conditioning and resistance to therapy.

Although this study establishes an important role for YB-1-positive sEVs in TNBC progression, several important questions remain unanswered. First, the molecular mechanisms governing selective loading of YB-1 into sEVs remain unclear. YB-1 is a multifunctional RNA- and DNA-binding protein (reviewed in [23]), raising the possibility that specific post-translational modifications, RNA interactions, or adaptor proteins facilitate its sorting into sEVs. Future studies should define the molecular machinery responsible for YB-1 packaging into sEVs and determine whether phosphorylation at S102 directly influences sEV incorporation. Second, the precise transcriptional programs induced by EV-delivered YB-1 in recipient cells remain to be determined. Genome-wide transcriptomic and epigenomic analyses will be important to identify the downstream targets activated following nuclear delivery of YB-1-positive sEV cargo. Such studies may reveal pathways involved in stemness, immune evasion, metastatic colonization, and therapeutic resistance. Third, while our work focused primarily on tumor cell-autonomous effects, the impact of YB-1-positive sEVs on stromal and immune cells within the tumor microenvironment and the pre-metastatic niche remain unknown. Because Vesiclepedia analysis identified YB-1-containing EVs in fibroblasts and immune cells, and YB-1 has been shown to regulate the immune microenvironment to enhance anti-tumor immune evasion [44, 49–54], it is plausible that tumor-derived YB-1-positive sEVs modulate macrophages, fibroblasts, endothelial cells, or myeloid populations to create a pro-metastatic microenvironment. Our recently published studies [15] have established a function of cancer-derived sEVs in activating tumor-associated fibroblasts. Future *in vivo* studies should investigate how YB-1-positive sEVs influence immune suppression, angiogenesis, extracellular matrix remodeling, and pre-metastatic niche formation.

The therapeutic implications of targeting YB-1 nuclear trafficking also warrant further investigation. Although PRR851 demonstrated potent anti-oncogenic effects *in vitro*, future studies are needed to evaluate its efficacy *in vivo* using orthotopic and metastatic TNBC models. It will also be important to determine whether inhibition of spathasome-mediated transport sensitizes tumors to chemotherapy, immunotherapy, or targeted therapies. Because YB-1 is implicated in drug resistance pathways [3], combined therapeutic strategies targeting both YB-1 signaling and conventional treatments may provide synergistic benefit. Additionally, it should be considered that the PRR851 mechanism of action is directed toward the whole sEV cargo, so that in addition to YB-1 the inhibition of nuclear translocation of additional proteins and/or RNA may have a compounding effect for the anti-oncogenic effect.

In summary, our study has identified YB-1-positive sEVs and YB-1 phosphorylation-dependent nuclear trafficking as critical regulators of intercellular oncogenic communication in TNBC. These findings uncover a novel mechanism by which EV-mediated nuclear signaling promotes malignant progression and suggest that targeting YB-1 transport pathways may represent a promising therapeutic strategy for aggressive breast cancers.

## Supporting information

Graphic Abstract

## Acknowledgements

This work was supported in part by grants R01CA272621 to KSA and U01CA304453 to KSA and AL.

## Declaration of Interest Statement

The authors declare a patent and pending patent applications entitled “Inhibition of a tripartite VOR protein complex in multicellular organisms” (EP3864409A1/US12527789B2), and “Use of triazole analogues for inhibition of the tripartite VOR complex in multicellular organisms” (EP4210696A1/US20230321082A1). The applicants and inventors include AL and MS, Touro University Nevada.

## Notes

### Competing Interest Statement

The authors have declared no competing interest.

