## Supplementary figures and images for "Nuclear translocation of phosphorylated YB-1 via small extracellular vesicles contributes to the malignant phenotype of triple negative breast cancer"

### Graphic Abstract

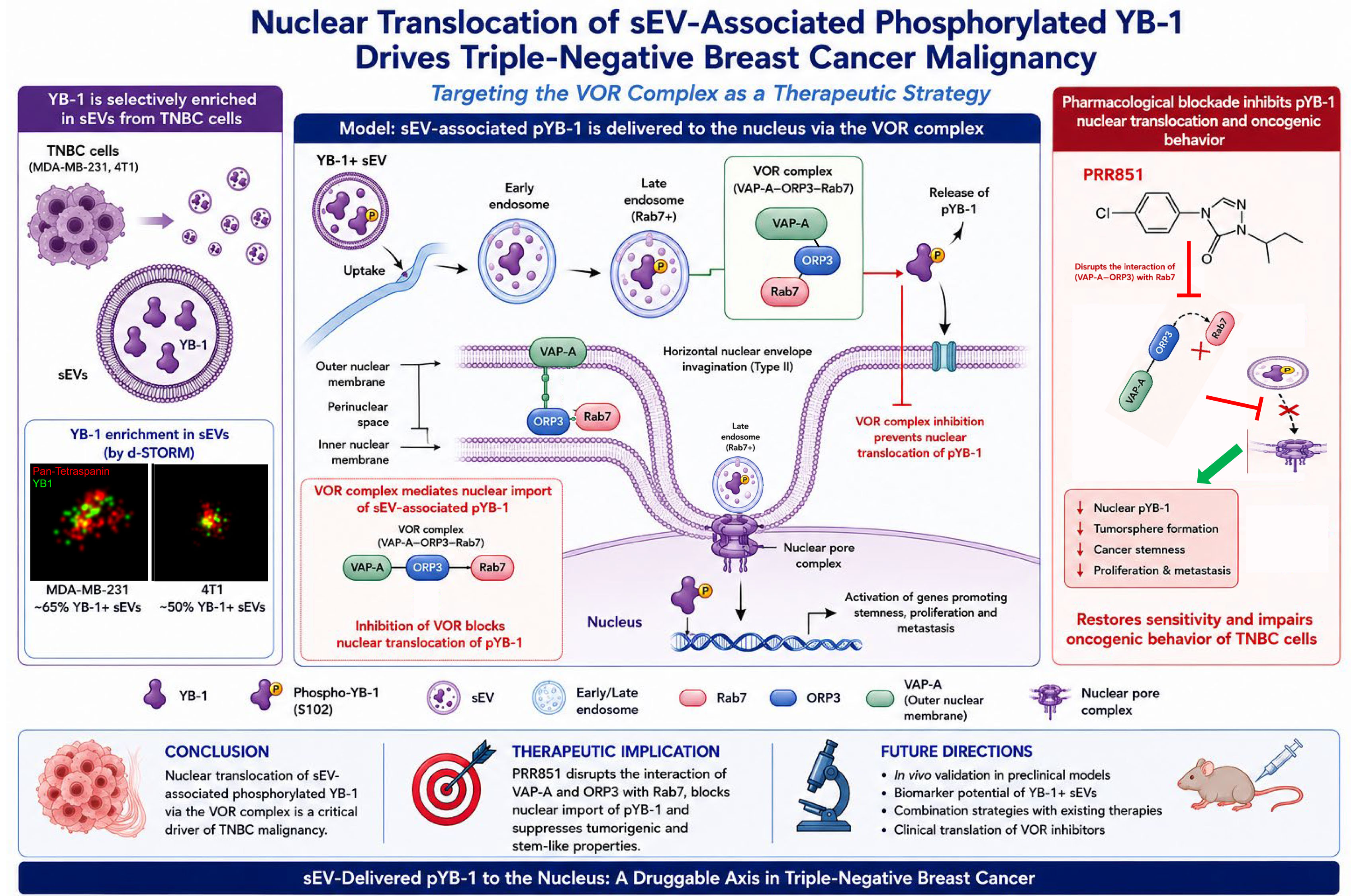

### Supplemental Table 1

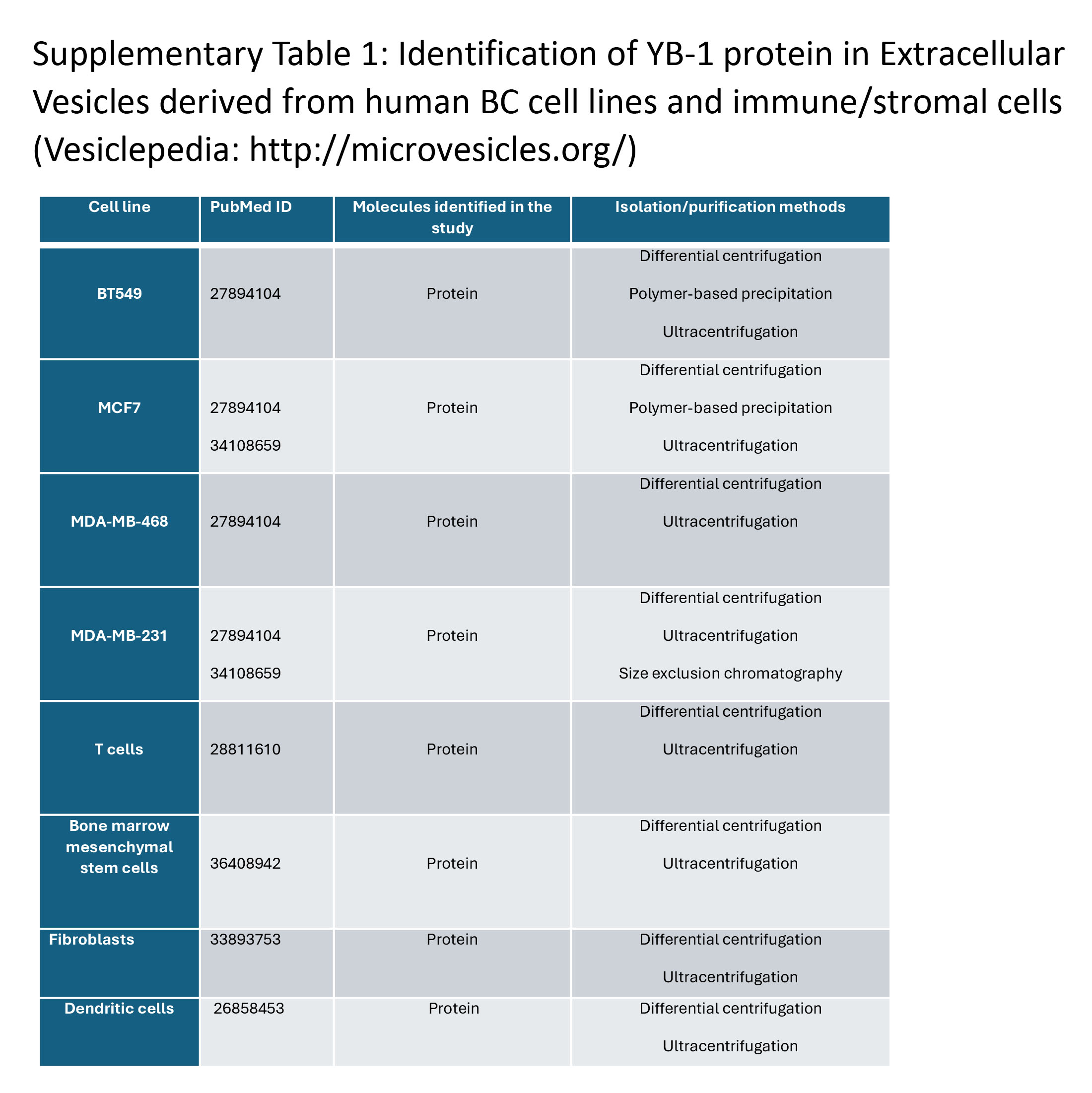
